# An improved culture protocol for the differentiation and survival of human promyelocytic leukaemia PLB-985 cell-line into mature neutrophil-like granulocytes

**DOI:** 10.1101/2021.09.14.460267

**Authors:** Shehu Shaayau, Andrew L. Cross, Helen L. Wright, Steven W. Edwards

**Affiliations:** Department of Biochemistry, Usmanu Danfodiyo University, Sokoto, Nigeria; Institute of Life Course and Medical Sciences, University of Liverpool, United Kingdom; Institute of Infection, Veterinary and Ecological Sciences, University of Liverpool, United Kingdom

**Keywords:** Cell culture, neutrophil, differentiation, PLB-985 cells, apoptosis

## Abstract

Circulating blood neutrophils are short-lived, lack proliferation capacity and cannot be transfected *in vitro* to express exogenous genes or proteins. These properties have made the *ex vivo* genetic manipulation of neutrophils challenging and hindered biochemical and molecular studies investigating the function of specific genes and proteins. Improved methodology for differentiating cell lines into mature neutrophil-like phenotypes, with similar morphological and functional properties to blood neutrophils would, therefore, be an important tool to probe the molecular properties of mature cells. The PLB-985 cell line was cultured in RPMI-1640 medium supplemented foetal calf serum (FCS) and penicillin/streptomycin. For induction of differentiation into neutrophil-like cells, the medium was supplemented with sodium pyruvate, *N, N*-dimethylformamide (DMF) and a*ll-trans* retinoic acid (ATRA), FCS and penicillin/streptomycin. The cytokines G-CSF and GM-CSF were used to enhance differentiation, prolong viability and delay the progression of the differentiated cells into apoptosis. The modified culture protocol and conditions induced PLB-985 cells to differentiate into mature, neutrophil-like granulocytes that resembled the morphology of mature blood neutrophils as evident by acquisition of a multi-lobed nucleus and granulated cytoplasm. These modified culture conditions resulted in enhanced differentiation into neutrophil-like cells and the apoptosis of these differentiated cells was delayed by supplementation with cytokines. This experimental system should be useful for studies probing the function of specific genes and proteins in human neutrophils.

## 1. INTRODUCTION

Polymorphonuclear leukocytes (PMNs) or neutrophils are key mediators of inflammation and play a major role in the body’s defence against microbial infection(Wright et al., 2010, Wright et al., 2014). They are the most abundant leukocytes in human blood and the first to be recruited to the site of infection or injury. Circulating blood neutrophils are often considered to be terminally differentiated, although they can be stimulated under appropriate conditions to up-regulate expression of sets of genes that alter their function and/or act as extracellular regulators of immune function (Cassatella et al., 2007, Wright et al., 2013, Thomas et al., 2015)). They are short-lived cells, lack proliferation capacity and rapidly undergo apoptosis following isolation from the blood (Akgul et al., 2001, Moulding et al., 2001). Because of these properties, they are extremely difficult to transfect *in vitro* and there are no published methods that describe reliable and reproducible protocols that enable expression of exogenous genes or proteins, despite many attempts to do so. Hence, this inability to genetically manipulate expression of key genes in human neutrophils has hampered studies aimed at characterising the molecular processes regulating infection or inflammation.

One approach that has been adopted by many research teams aiming to genetically manipulate neutrophil gene expression has been to use culturable myeloid cell lines capable of differentiating into mature neutrophil-like cells *in vitro*, to acquire the molecular and functional properties of mature neutrophils (Collins et al., 1978, Pivot-Pajot et al., 2010, Tucker et al., 1987). Many such myeloid cell lines have been used, and there have been some successes in this approach. For example, cell lines such as HL-60 and PLB 985 have been transfected *in vitro* to alter expression of key genes and then treated with agents such as retinoic acid, to induce differentiation into more mature neutrophil-like cells (Witko-Sarsat et al., 2010, Moriceau et al., 2009, Ashkenazi and Marks, 2009, Pessach and Levy, 2000). However, while such systems may well be capable of altering gene expression, the efficiency of such systems is very mixed in that (a) there is highly-variable differentiation into neutrophil-like cells and acquisition of landmark features such as a multi-lobed nucleus and expression of key surface receptors and (b) mature neutrophils spontaneously undergo apoptosis and hence the lifespan of these differentiated cells is very low.

The human promyelocytic leukaemia cell line, PLB-985 is a cell line that was originally described as having been established from the peripheral blood of a patient with acute non-lymphocytic leukaemia (Tucker et al., 1987) but later shown to be a sub-clone of HL-60 (Drexler et al., 2003). PLB-985 cells have been reported to show higher levels of differentiation into neutrophil-like granulocytes upon appropriate culture (Pedruzzi et al., 2002). The cell line has been shown to undergo differentiation into both granulocytes and monocytes/macrophages in the presence of appropriate inducing agents (Sanmun et al., 2009, Pedruzzi et al., 2002). However, the efficiency of differentiation that has been reported is variable (and often very low), and the morphology of the differentiated cells only partly resembles that of mature neutrophils. In addition, mature neutrophils constitutively undergo apoptosis (Fox et al., 2010) and so the more efficient the differentiation of PLB-985 cells into mature neutrophils, the greater the extent of apoptosis. This enhanced rate of apoptosis of differentiated cells hampers their usefulness as experimental models.

The aim of the present study was two-fold. First, we aimed to develop a more efficient culture methodology for the differentiation of PLB-985 cells into neutrophil-like cells. In particular, we wanted to devise a culture protocol that led to maximal efficiency of differentiation of cells with a multi-lobed nucleus, a hallmark feature of mature neutrophils. Secondly, because mature neutrophils spontaneously undergo apoptosis during culture *in vitro*, we aimed to modify the culture/differentiation conditions to maximally enhance the survival kinetics of the differentiated cells.

## 2. MATERIALS AND METHODS

The promyelocytic leukaemia PLB-985 cell line was a kind gift from Dr Veronique Witko-Sarsat (Cochin Institute, Paris). RPMI 1640 (+10mM L-glutamine), RPMI 1640 + HEPES (+10mM L-glutamine), foetal calf serum (FCS), *all trans* retinoic acid (ATRA) (R2625), *N,N*-dimethyl formamide (DMF), sodium pyruvate (100 mM), and all other reagents were from Sigma-Aldrich (Dorset, UK).

### 2.1 Culture and differentiation of PLB-985 cells

PLB-985 cells were routinely cultured in RPMI-1640 (+10mM L-glutamine) medium (supplemented with 10% (v/v) foetal calf serum and 1% (v/v) penicillin/streptomycin) and incubated at 37°C in a humidified atmosphere of 5% CO_2_. Cell cultures were passaged every 2 - 3 days or as indicated in the text. For induction of differentiation into granulocytes (Table 1), RPMI-1640 (+10mM L-glutamine) medium was supplemented with 0.5% (v/v) foetal calf serum, 1% (v/v) penicillin/streptomycin, 1 mM sodium pyruvate, 0.5% (v/v) *N, N*-dimethylformamide (DMF) and 1µM *All trans* retinoic acid (ATRA). Unless stated in the text, the culture was passaged and re-suspended in fresh medium on days 2 and 4. Exponentially growing PLB-985 cells were seeded at a starting density of 2×10^5^/mL and defined as dPLB-985 (differentiation-induced) and PLB-985 (non-induced) cells. Incubations were for up to a 6/7-day incubation period.

**Table 1.** Composition of culture media.

| Constituent | Maintenance medium | Differentiation medium |
| --- | --- | --- |
| RPMI 1640 + 10mM L-glutamine. | 500 mL | 500 mL |
| Foetal calf serum (FCS) | 10% (v/v) | 0.5% (v/v) |
| Penicillin/Streptomycin | 1% (v/v) | 1% (v/v) |
| Sodium pyruvate | - | 1 mM |
| *N, N*-dimethylformamide (DMF) | - | 0.5% (v/v) |
| *All-trans* retinoic acid (ATRA) | - | 1 μM |

Prior to analysis, cells were centrifuged at 1,000 g for 3 min and the supernatant discarded. Cells were re-suspended in fresh media, counted using a Multisizer3 cell counter (Beckman Coulter) and the suspension volume adjusted with media to give a final concentration of 1×10^6^/mL.

### 2.2 Isolation of human blood neutrophils

Ethical approval was obtained for this study from the University of Liverpool Committee on Research Ethics (CORE). All participants gave written, informed consent. Whole blood was collected from healthy donors by venipuncture, into heparinised vacutainers and polymorphonuclear leukocytes were separated by centrifugation using Ficoll-paque according to manufacturer’s instructions (Wright et al., 2017). Whole blood was mixed with HetaSep solution at a ratio of 1:5 (HetaSep:whole blood) and incubated at 37°C for 30 min until the plasma/erythrocyte interphase was at ∼50% of the total volume. The leukocyte-rich plasma layer was carefully removed and diluted in a 4-fold volume of media: Mg^2+^- and Ca^2+^-free PBS, 2% FBS, and 1 mM EDTA). Cells were centrifuged at 200 *g* for 10 min and resuspended in a 4-fold volume of the same media. Washed leukocytes were layered onto Ficoll-Paque 1:1 and centrifuged at 500 *g* for 30 min. After centrifugation, the cell pellet was re-suspended in RPMI-1640 + HEPES (10mM L-glutamine) media and then contaminating erythrocytes removed by hypotonic lysis using ammonium chloride buffer (155 mM NH_4_Cl, 13.4 mM KHCO_3_, 96.7 μM EDTA, PH 7.6), in a 1:9 ratio of cell media to lysis solution, for 3 min followed by centrifugation at 1,000 g for 5 min. Cells were suspended in RPMI-1640 + HEPES (10mM L-glutamine) media and final concentration adjusted to 5 ×10^6^ cells/mL using RPMI-1640 + HEPES (10mM L-glutamine) medium. Cell purity was assessed after cytospins (10^5^ cells in 200 μL PBS) at 500 g for 5 min in a Shandon3 cytocentrifuge, and routinely found to be >95%, as determined by morphological examination under a light microscope, after Rapid Romanowsky staining. Cell viability immediately after isolation, measured by Guava viaCount on a Guava EasyCyte instrument, was also routinely >95%.

### 2.3 Morphological analysis of PLB-985 cells

Following culture of PLB-985 cells in the presence and absence of differentiation-inducing agents, 20μL of a cell suspension (at a concentration of 1×10^6^/mL) was diluted with 180μL PBS (EDTA), and slides prepared using a Shandon3 cytospin after cytocentrifugation at 500 × g for 5 min to fix the cells on the slides. The slides were air-dried, stained with Rapid Romanowsky stain and cell morphology analysed under a light microscope. Cells with a large, rounded nucleus with well-defined nucleoli and agranular cytoplasm were observed in non-differentiated PLB-985 cells while the differentiated cells appeared with segmented, multi-lobed nuclei and granular cytoplasm.

### 2.4 Statistics

Statistical analysis was carried out using SPSS v24, using Student’s t-test unless otherwise stated.

## 3. RESULTS

### 3.1. Differentiation conditions 1

In initial experiments, PLB-985 cells were cultured under two experimental conditions as shown in Table 1, defined as PLB-985 (maintenance medium) and dPLB-985 (differentiation medium). Cells cultured in PLB-985 and dPLB-985 both showed an exponential growth pattern with a doubling time of approximately 24h (Figure 1A) during first few days of the culture (days 1-3). Thereafter, the proliferation rate declined as the cells aged in the culture. By day 3, the cell number was lower in the dPLB-985 cultures due to the loss of proliferation capacity caused by ATRA, DMF and sodium pyruvate. However, as the cell culture period extended beyond 3 days, the rate of cell proliferation declined, and this was most marked for the dPLB-985 cells. In both cell cultures numbers declined by days 6 and 7 (Figure 1A).

**Figure 1.**
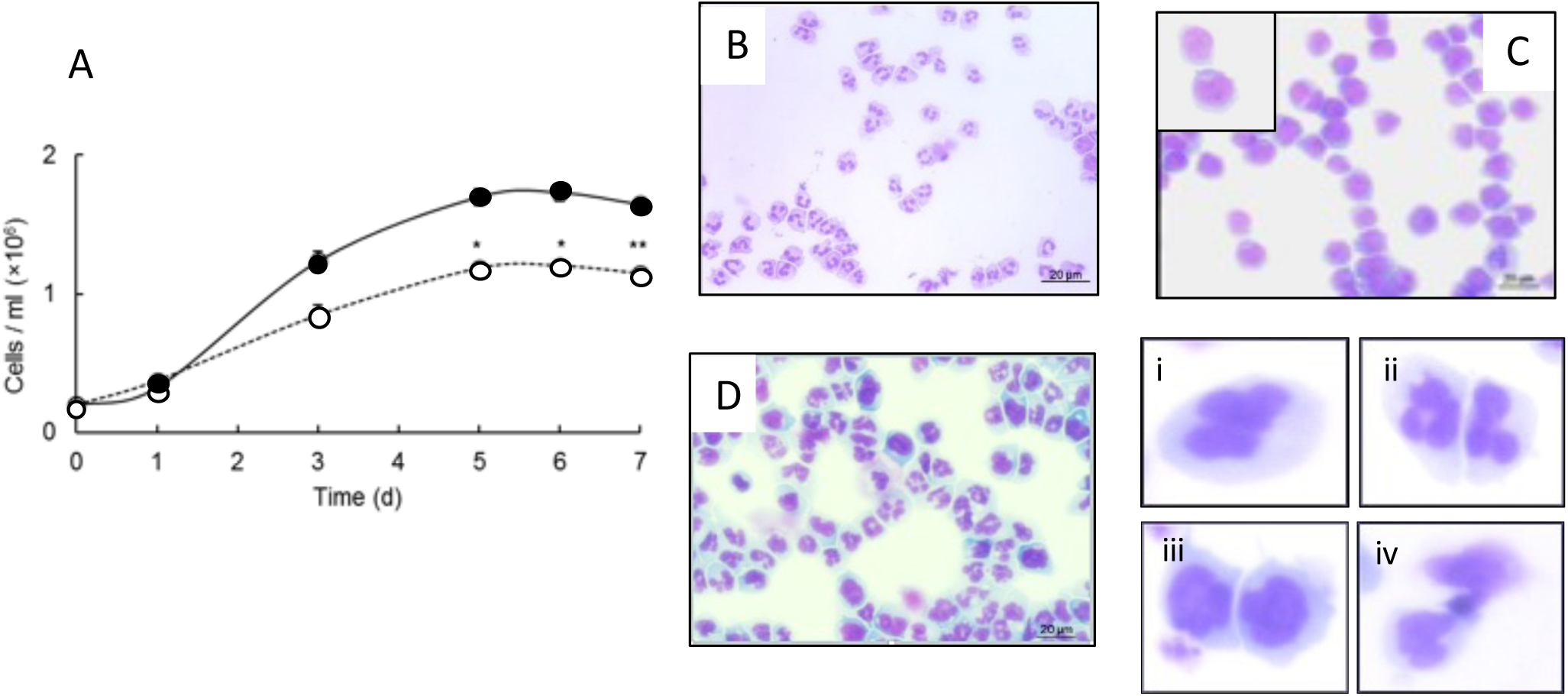
Growth of PLB-985 cells in the absence and presence of differentiation-inducing agents. In A, exponentially growing PLB-985 cells at a starting density of 2×10^5^/mL were cultured in separate flasks designated PLB-985 (non-differentiating •) and dPLB-985 (differentiation-induced ○) cells, as described in Table 1. The media was unchanged for 7 days and cell numbers were counted on days 0, 3, 5, 6 and 7 using a Multisizer 3 coulter counter, following 1:1000 dilution with Isoton II. In non-differentiating media, the cells had a doubling time of 24 h initially, but after day 3, the growth rate decreased, particularly in dPLB culture. Data are expressed as mean of total cells (± SEM, n=3), * = p≤0.05, ** = p≤0.01 (paired, two-tailed student’s t-test). B shows cytospin of purified human neutrophils, showing their characteristic multi-lobed nucleus and granular cytoplasm, while C, shows cytospin of non-differentiated PLB cells which were larger than human neutrophils and had a large, single-lobed nucleus that largely filled the cytoplasm. D shows a cytospin of dPLB-985 cells after 5 days of culture, with many cells showing signs of differentiation into neutrophil-like cells with multi-lobed nuclei and granular cytoplasm. Enlarge images show (i) partially-differentiated cells, (ii) fully-differentiated cells, (iii) undifferentiated cells and (iv) apoptotic cells with small, condensed nuclei.

At time points throughout this culture, samples were removed for analysis of cell morphology by Romanowsky staining of cytospin slides and light microscopic examination to determine cell viability and morphology. Differentiation was assessed by morphological criteria and compared to the morphology of mature, blood neutrophils (Figure 1B). These cells are characterised by their relatively small cell size (∼8-10 µm diameter) a multi-lobed nucleus and granular cytoplasm. These features were used to compare the morphological changes associated with differentiation of PLB-985 cells.

PLB-985 cells cultured in non-differentiation medium (PLB-985) had a round, diffuse nucleus that occupied a large proportion of the cytoplasm (Figure 1C) and no morphological features associated with differentiation were observed at any time points during the 7-day culture. When PLB-985 cells were cultured in differentiation medium (dPLB-985), they began to show signs of differentiation, such as indentations and convolutions in the nuclear membrane after day 2 of differentiation, and by day 5-6, many cells had acquired morphological features of a mature neutrophil, such as decreased cell size, segmentation of the nuclei into 3-5 lobes, disappearance or decreased appearance of nucleoli and granulation of the cytoplasm (Figure 1D). Some cells showed changes in nuclear morphology, but in some, the nucleus could not be termed multi-lobed (Figure 1.i). Instead, these cells resembled band cells (not fully mature neutrophils) and hence, such cells are defined as “partially differentiated”. Some cells showed clearly defined multi-lobed nuclei (Figure 1.ii), and hence these were defined as “differentiated cells”. Some cells, however, showed few, if any signs of differentiation as assessed by nuclear morphology (Figure 1.iii) and were termed “non-differentiated cells”. Cells with a condensed nucleus were defined as apoptotic cells (Figure 1. iv). These morphological changes were then quantified as the cells were cultured for up to 6 days in either medium. In the dPLB-985 medium there was progressive differentiation towards neutrophil-like cells which was evident from day 3 in culture and by day 5, >75% of the cells (Table 2) showed signs of differentiation, with ∼65% having a fully defined, multi-lobed nucleus, a characteristic of mature neutrophils. In parallel, the number of non-differentiated cells declined. After day 3, a mixture of different cell populations was observed in the dPLB-985 medium, resulting from various stages of differentiation undergone by the cells (Figure 2) that included undifferentiated cells, partially mature band-like cells, differentiated cells and apoptotic/dead cells. By Day 6 and beyond, dead/apoptotic cells accounted for >20% of the population, which were visible as small, round cells with a condensed nucleus. Also, by Day 7, the total number of cells had decreased, presumably as a result of lysis of the apoptotic cells. While the above experiments indicate that the differentiation medium dPLB-985 could successfully differentiate cells into neutrophil-like cells with the characteristic multi-lobed nucleus, the number of apoptotic/dead cells increased over time in culture. This is likely explained by the fact that mature neutrophils have a very short half-life in culture (Moulding et al., 2001) and rapidly undergo apoptosis. Therefore, this culture and differentiation protocol was modified with the aims of trying to (a) increase the percentage of fully differentiated cells and (b) to decrease the rate of apoptosis once the cells had differentiated into mature cells.

**Table 2.** Proportions of dPLB-985 cells after day 3-6 of induction. PLB-985 cells with different morphologies based on their level of differentiation (See Figure 1), counted manually by morphological examination of cells from representative cytospin slides viewed under the light microscope. Counts are expressed as percentages of the total number of cells (± SD, n=3).

| Cells Differentiation Status | Day 3 | Day 5 | Day 6 |
| --- | --- | --- | --- |
| Differentiated (%) | 44.41 ± 4.07 | 65.89 ± 5.17 | 59.33 ± 4.39 |
| Partially differentiated (%) | 24.74 ± 4.32 | 13.35 ± 2.04 | 10.94 ± 4.24 |
| Non-differentiated (%) | 26.44 ± 3.54 | 8.41 ± 4.16 | 6.21 ± 3.36 |
| Apoptotic (%) | 4.41 ± 1.36 | 12.35 ± 3.61 | 23.52 ± 4.43 |
| Total | 100 | 100 | 100 |

**Figure 2.**
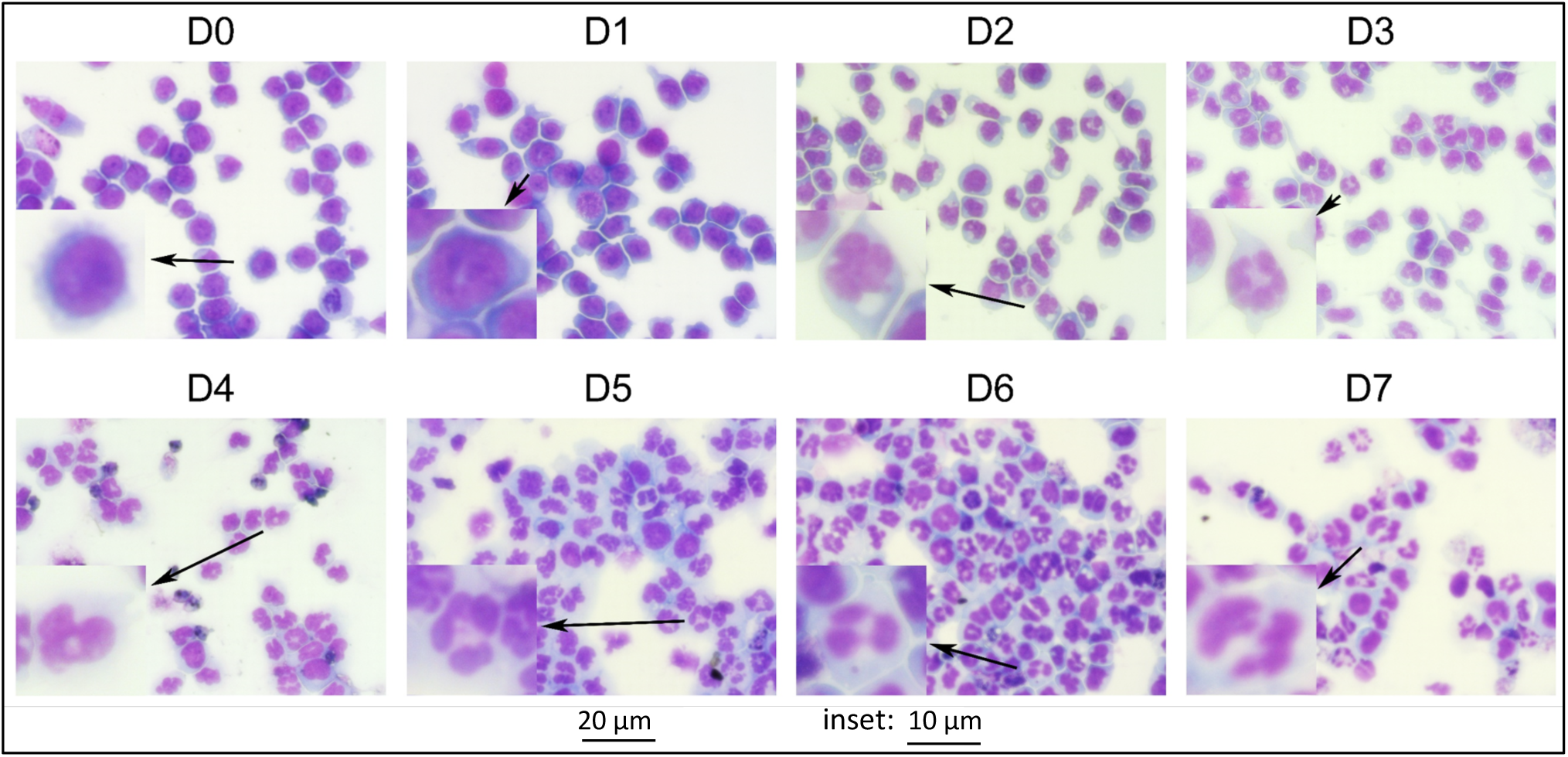
Morphology of dPLB-985 cells during 7 days culture. dPLB-985 cells were cultured in the presence of differentiation-inducing agents for 7 days and cell samples were removed daily, cytospin slides prepared, stained with Romanowsky stain and viewed under the light microscope. Large, non-lobular nuclei, frequent nucleoli and agranular cytoplasm were evident on day 0-1. After day 2 of induction, cells began to show signs of differentiation, such as indentations in the nucleus, decrease in number of nucleoli, and appearance of granules in the cytoplasm. By days 4-7, the cells had acquired multi-lobular nuclear morphology. Some cells exhibited apoptotic morphologies from day 4.

### 3.2. Differentiation conditions 2

The first modification to the differentiation protocol was to replace the media on days 2 and 4 of the incubation period. The aim of this was to remove any factors that could inhibit growth or differentiation or promote apoptosis, (e.g. pH, toxic products) and/or to replenish any factors that became limiting. Replacement of media on days 2 and 4 of the culture period, resulted in a significantly increased percentage of differentiated cells, compared to cells that were cultured for 7 days continuously in unchanged medium (Figure 3A). For example, in this series of experiments, the number of differentiated cells after 5 days of culture in unchanged medium was 60%, which increased to 72% (p<0.01) when the medium was preplaced at days 2 and 4. In these experiments, cell viability was also measured using the Guava Viacount assay which measures changes in the permeability of the plasma membrane. In the absence of changes in the culture medium at days 2 and 4, the viability of the differentiated cells, which was close to 100% over the first two days of culture, which gradually decreased to around 20% by day 5 and then decreased further by day 7 (Figure 3B). However, in differentiated cultures in which the media was replaced with fresh media at days 2 and 4, there was a statistically-significant (p<0.01, n=3) increase in the viability from day 3 onwards, compared to cells without media changes. By day 5, ∼60% of the cells were viable after the media changes, whereas only ∼20% were viable in the absence of these media changes.

**Figure 3.**
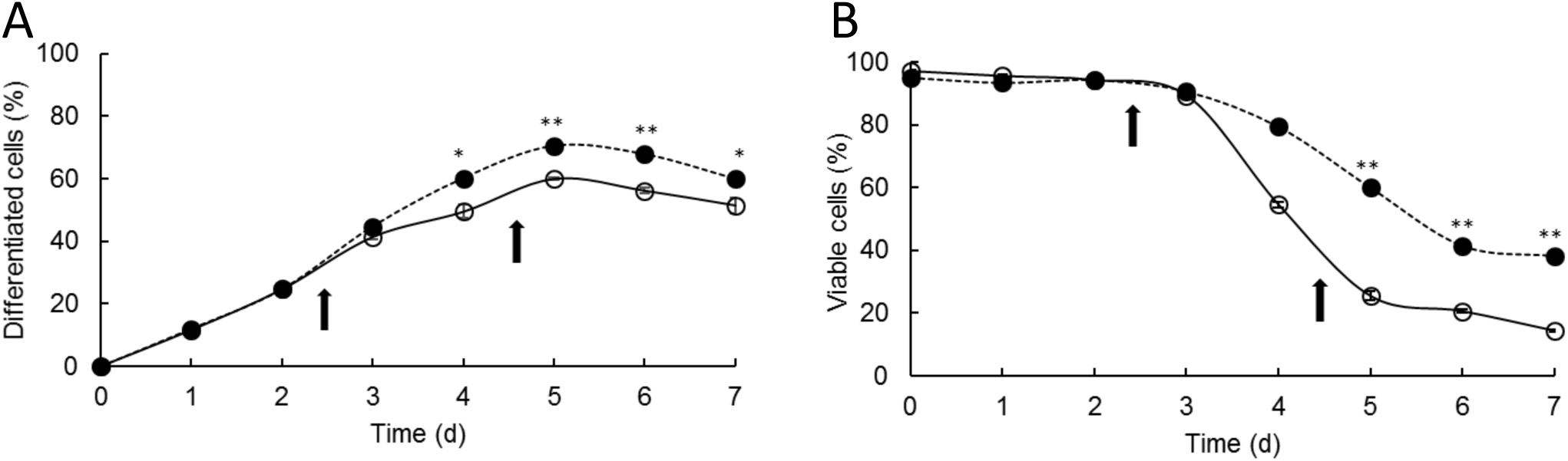
Effect of media changes on differentiation and viability of dPLB-985 cells. PLB-985 cells were cultured in differentiation media for 7 days, without (○) and with media changes on days 2 and 4 (•). The number of differentiated cells was counted manually from representative cytospin slides viewed under the light microscope and the viability was measured using the viaCount assay on the flow cytometer. (A) Shows the number of differentiated PLB-985 cells and (B) shows dPLB-985 cell viability. Arrows indicate times of media changes and data are expressed as percentage of total cells (± SEM, n=3), * = p≤0.05, ** = p≤0.01 (paired, two-tailed student’s t-test).

### 3.3 Differentiation conditions 3

A further modification of the differentiation procedure was then performed by modifying the improved differentiation conditions described above (changing the medium at days 2 and 4) but supplementing the differentiation media with agents known to promote granulopoiesis and/or delay apoptosis in mature blood neutrophils (Burgess and Metcalf, 1980, Souza et al., 1986, Begley et al., 1987, Derouet et al., 2006). Hence, the pro-inflammatory cytokines and granulocyte maturation agents, G-CSF (10 ng/mL) and GM-CSF (5 ng/mL) were added to the differentiation culture, to test their effects on differentiation and viability of the differentiated PLB-985 cells. These complete media were replaced after days 2 and 4 of culture.

As shown in Figure 4A and B, the addition of these two cytokines further increased the number of differentiated cells. For example, by day 5 G-CSF significantly (p≤0.05, n=3) increased the proportion of differentiated cells from 70.5% ± 0.4% (with media replacements only) to 79.4% ± 1.9% (Figure 4A). GM-CSF also significantly increased (p≤0.05, n=3) the proportion of differentiated cells to 81.7% ± 1.1%, (Figure 4B). Whilst these small increases in the numbers of differentiated cells were significant, more dramatic effects were observed on cell viability. For example, after 7 days in culture, cell viabilities were increased from 38.2% ± 0.8% (with media changes only) to 66.8% ± 0.9% by G-CSF (Figure 5A) and to 68.1% ± 4.5% for GM-CSF (Figure 5B) (p≤0.01, n=3).

**Figure 4.**
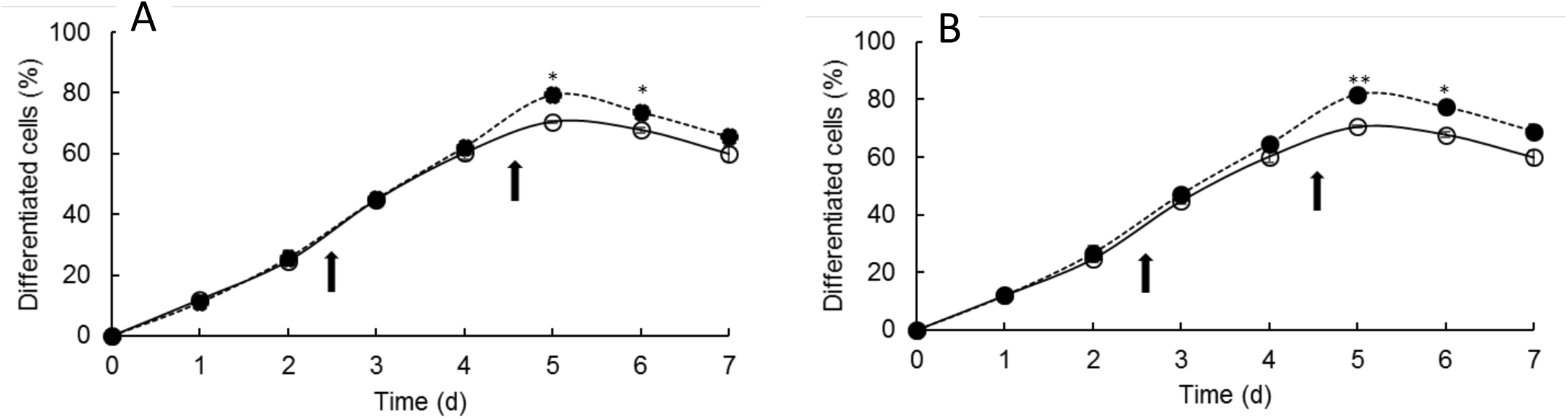
Effects of G-CSF and GM-CSF on differentiation of PLB-985 cells. dPLB-985 cells were incubated without (○) and with (•) addition of cytokines G-CSF, 10ng/mL (A) or GM-CSF, 5ng/mL (B) after day 2 and 4 following media changes. The number of differentiated cells was counted manually from morphological examination of representative cytospin slides viewed under the light microscope. Arrows indicate times of media changes and cytokine supplementation (after days 2 and 4). Data are expressed as percentages of total cells. (± SEM, n=3), * = p≤0.05, ** = p≤0.01 (paired, two-tailed student’s t-test).

**Figure 5.**
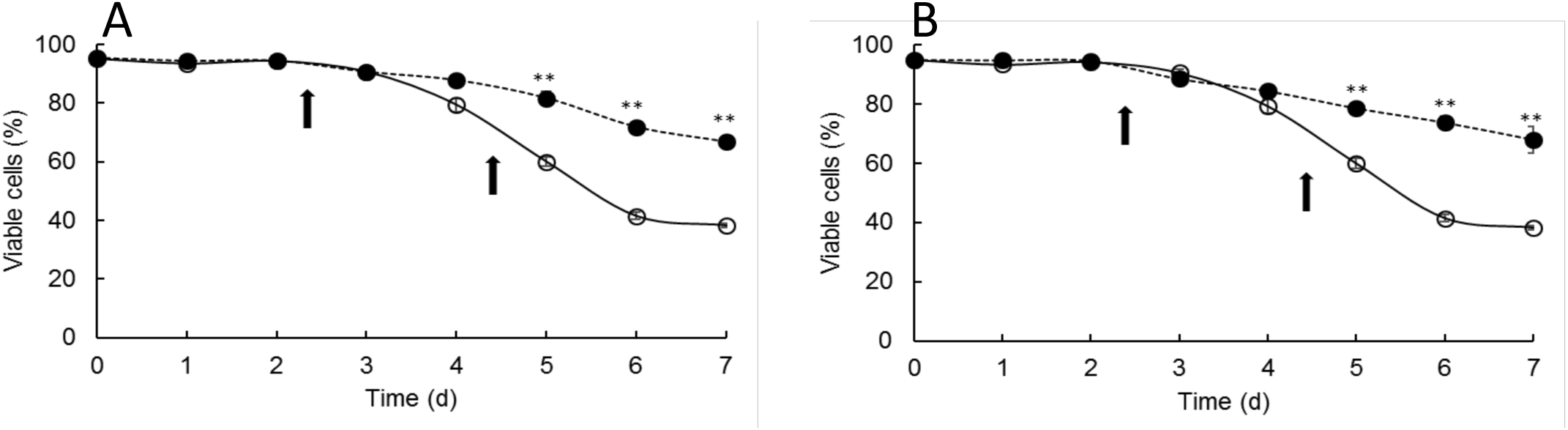
Effects of G-CSF and GM-CSF on viability of dPLB-985 cells. Differentiating PLB-985 cells were incubated without (○) and with (•) addition of cytokines G-CSF, 10ng/mL (A) or GM-CSF, 5ng/mL (B) after day 2 and 4 following media changes. The cell viability was measured using the viaCount assay on the flow cytometer. Arrows indicate times of media changes and cytokine supplementation (after days 2 and 4). Data are expressed as percentages of total cells. (± SEM, n=3), * = p≤0.05, ** = p≤0.01 (paired, two-tailed student’s t-test).

## 4. DISCUSSION

The human promyelocytic leukaemia PLB-985 cell line has the capacity to undergo both granulocyte and monocyte/macrophage differentiation in the presence of appropriate chemical inducing agents. Phenotypically, they are myelomonoblast-like cells, and upon exposure to the appropriate agents, they can be stimulated to differentiate into mature granulocytes or monocytes/macrophages (Tucker et al., 1987). This study identified the culture conditions that induced differentiation of PLB-985 cells into mature neutrophil-like phenotypes.

The assessment of differentiation in this study was mainly based on morphological properties. Standard morphologic/staining criteria categorizes myeloid cells into: immature; comprising blasts, promyelocytes and promonocytes, or mature; comprising myelocytes, metamyelocytes, bands and segmented neutrophils (Chomienne et al., 1990). The non-differentiated PLB-985 cells are characterised as being at the promyelocytic stage, but the combined effects of ATRA, DMF and sodium pyruvate have successfully transformed them along the myeloid lineage into mature cells identified as band neutrophils and segmented neutrophils, the final cells in the myeloid lineage. This pattern of differentiation is analogous to that seen in neutrophil differentiation and maturation (Iwasaki and Akashi, 2007). The differentiated PLB-985 cells exhibited other morphological features of mature neutrophils, including reduced cell size, indented, convoluted and segmented nuclei, decreased number or absence of nucleoli and granulated cytoplasm (Iwasaki and Akashi, 2007). After 3-4 days of induction, metamyelocytes and band neutrophils were the predominant cells in the culture in differentiation medium, whilst after 5-7 days in culture, segmented neutrophils predominated. In contrast, PLB-985 cells cultured in the absence of differentiation-inducing agents remained in their promyelocytic form, exhibiting large, rounded nuclei containing 2-4 nucleoli each, with a dispersed chromatin, agranular cytoplasm and relatively high nuclear/cytoplasmic ratio.

Interestingly, alongside the progressive increase in proportion of cells showing mature neutrophil morphology, there was a parallel increase in the proportion of cells showing morphological features of apoptosis, such as condensation of chromatin and loss of nuclear lobes (Dransfield et al., 1994). Indeed, these apoptotic cells only appeared in appreciable numbers after the differentiated PLB-985 cells when mature neutrophil morphology predominated in the culture, usually after day 5-7 of culture. Apoptosis is a non-pathological, highly-programmed cell death that is activated constitutively when neutrophils age *in vivo* and *in vitro* (Kennedy and DeLeo, 2009). Thus, the differentiated PLB-985 cells were considered to have acquired one of the most definitive features of mature blood neutrophils; namely spontaneous progression into apoptosis. These observations confirm that the differentiated PLB-985 cells may be a good cell line model for the *in vitro* study of neutrophil signalling and functions, such as the underlying mechanisms of neutrophil apoptosis.

The measurement of PLB-985 cells growth rate following induction of differentiation demonstrated that differentiation significantly decreased the rate of cell growth due to the loss of proliferation capacity caused by ATRA (Chomienne et al., 1990, Degos and Wang, 2001) and DMF (Pedruzzi et al., 2002). Therefore, as the cells began to differentiate, their proliferation capacity declined and when they became terminally differentiated, they ceased to proliferate and then undergo apoptosis constitutively, thus behaving more like mature blood neutrophils. Hence, the new differentiation protocol described in this study has proved effective in inducing PLB-985 cells to differentiate into mature neutrophil-like granulocytes with the typical multi-lobed nucleus and granulated cytoplasm.

The effects of differentiation media changes on days 2 and 4 of culture demonstrated that the proportion of differentiated PLB-985 cells increased significantly as a result of the media replacements during the culture period. Most likely, these changes of fresh medium replenished the nutrient supply to the differentiating cells, kept the appropriate pH of the culture and eliminated the accumulated metabolic wastes from it. The media changes caused a slight increase in the viability of the differentiated PLB-985 cells as well. There was, however, a drastic decrease in the number of viable cells that was evident after day 4 of culture, which is attributable to the constitutive apoptosis within around 24h of the differentiation.

Another significant finding in this study was the importance of cytokines G-CSF and GM-CSF supplementation in enhancing both the efficiency of differentiation and viability of the differentiated PLB-985 cells. These observations correlated with the previously reported studies on the effects of G-CSF and GM-CSF on proliferation, differentiation and survival of granulocytes (Chiewchengchol et al., 2015, Derouet et al., 2004, Maianski et al., 2002). The induction of HL-60 cells to differentiate terminally into mature granulocytes through exposure to the two cytokines has been reported (Souza et al., 1986, Elias and Van Epps, 1988). G-CSF has been shown to stimulate the formation of neutrophil-like granulocyte colonies in semi-solid agar, induce the terminal differentiation and inhibit the self-renewal capacity of murine myelomonocytic leukaemia cells *in vitro* (Burgess and Metcalf, 1980). In a similar manner, GM-CSF has also been demonstrated to promote proliferation and function of myelomonocytic cells, increase the number of neutrophils by delaying apoptosis and induce neutrophil functions, such as priming of reactive oxygen generation (Edwards et al., 1989). Mature blood neutrophils undergo apoptosis constitutively and are therefore short-lived cells. However, this spontaneous apoptosis can be delayed and neutrophil lifespan considerably prolonged by exposure to the GM-CSF, which acts by increasing their levels of survival protein Mcl-1 (Derouet et al., 2004, Derouet et al., 2006, Moulding et al., 2001). Mcl-1 is an anti-apoptotic member of Bcl_-_2 family proteins which has a very high turnover rate, and whose cellular levels correlates with survival status of the neutrophil (Thomas et al., 2010). It has been shown that GM-CSF enhances the cellular levels of Mcl-1 by promoting its stability and reducing its turnover rate that occur normally via the proteasome (Derouet et al., 2004).

To our knowledge, the percentage and survival of differentiated neutrophil-like cells with a multi-lobed nucleus and cytoplasmic granulation described in this study, are far superior to other reports using this, or other cell myeloid cell lines. Furthermore, apart from this high efficiency of PLB-985 cell differentiation into mature neutrophil-like granulocytes, the optimised conditions used in this study have also greatly enhanced the lifespan of the differentiated PLB-985 cells.

In conclusion, we describe here an improved methodology to differentiate PLB-985 cells into cells with a morphology that is characteristic of mature human neutrophils. In addition, we describe culture conditions that extend the lifespan of these differentiated cells.

## 5. FUNDING

SS was funded by a Commonwealth PhD scholarship from the Commonwealth Scholarship Commission.

## Notes

### Competing Interest Statement

The authors have declared no competing interest.

